# Integrating end-to-end learning with deep geometrical potentials for *ab initio* RNA structure prediction

**DOI:** 10.1101/2022.12.30.522296

**Authors:** Yang Li, Chengxin Zhang, Chenjie Feng, Peter L. Freddolino, Yang Zhang

**Author notes:** Correspondence to Peter L. Freddolino and Yang Zhang.

## Abstract

RNAs are fundamental in living cells and perform critical functions determined by the tertiary architectures. However, accurate modeling of 3D RNA structure remains a challenging problem. Here we present a novel method, DRfold, to predict RNA tertiary structures by simultaneous learning of local frame rotations and geometric restraints from experimentally solved RNA structures, where the learned knowledge is converted into a hybrid energy potential to guide subsequent RNA structure constructions. The method significantly outperforms previous approaches by >75.6% in TM-score on a nonredundant dataset containing recently released structures. Detailed analyses showed that the major contribution to the improvements arise from the deep end-to-end learning supervised with the atom coordinates and the composite energy function integrating complementary information from geometry restraints and end-to-end learning models. The open-source DRfold program allows large-scale application of high-resolution RNA structure modeling and can be further improved with future release of RNA structure databases.

## Introduction

RNA molecules perform a broad range of important cellular functions, ranging from gene transcription, regulation of gene expression, scaffolding, to catalytic activities. The critical functional roles of RNAs make them a new type of drug targets. It was estimated that targeting RNAs with small molecules will expand the drug design landscape by more than an order of magnitude compared to traditional protein-targeted drug discovery (Warner et al., 2018). Since many of the biological functions depend on specific tertiary structures of the RNAs, it becomes imperative to determine 3D structures of RNAs, in order to facilitate RNA-based function annotations and drug discovery. The biophysical experiments capable of resolving RNA structures, e.g., X-ray crystallography, Cryogenic Electron Microscopy (Cryo-EM) and Nuclear Magnetic Resonance (NMR) Spectroscopy, are unfortunately cost- and labor-intensive. Fast and accurate computational approaches for sequence-based RNA structure modeling are therefore highly needed.

Traditional RNA structure prediction approaches are often designed to construct 3D RNA structures from homologous modeling and/or physics-based simulations. For examples, methods such as ModeRNA (Rother et al., 2011) and RNABuilder (Flores et al., 2010) extract structure information from previously solved structural templates. For RNA targets with divergent sequences or novel topologies, their performance is usually not satisfying. Another family of methods, typified by RNAComposer (Biesiada et al., 2016) and 3dRNA (Zhao et al., 2012), assembles full-length RNA structures from fragments searched from a prebuilt fragment library. *Ab initio* RNA structure prediction methods, such as SimRNA (Boniecki et al., 2016) and RNA-BRiQ (Xiong et al., 2021), apply statistical potentials to guide the structure folding simulations. Despite the advantage that they do not directly rely on templates, the accuracy of the fragment assembly and *ab initio* folding approaches is low in general.

Deep machine learning has recently demonstrated promising performance in RNA structure feature predictions. For example, SPOT-RNA (Singh et al., 2019) and e2efold (Chen et al., 2020) utilized convolutional neural network (CNN), recurrent neural network (RNN) or transformer techniques to improve the accuracy of secondary structure (SS) predictions of RNAs. Analogous to the highly successful efforts in protein contact and distance predictions (Li et al., 2019), SPOT-RNA-2D (Singh et al., 2022) and RNAcontact (Sun et al., 2021) applied deep residual networks to learn inter-nucleotide distance/contacts from profile covariance. Despite the interest in the property learning, few studies have performed full-length modeling of RNA tertiary structures. Recently, a deep equivariant model, ARES (Townshend Raphael et al., 2021), was trained to score the conformations from limited data. However, it requires sufficient pre-sampled conformations, which may limit the application.

Inspired by the recent success of deep learning techniques in 3D protein structure prediction (Baek et al., 2021; Jumper et al., 2021; Li et al., 2022), we proposed a new deep learning pipeline, DRfold, to improve the performance of *ab initio* RNA structure prediction. Different from the full-atom end-to-end training in protein (Jumper et al., 2021), we adopted a coarse-grained model of RNA specified by the phosphate P, ribose C4’ and glycosidic N atom of the nucleobase, for training efficiency. In particular, we added a geometric module which is trained in parallel to assist the end-to-end training, and meanwhile aggregates both end-to-end and geometric potentials to guide subsequent RNA structure reconstruction simulations. We found that the integration of the end-to-end training and deep geometric learning, followed by the gradient-based optimization, generates RNA structure models with accuracy significantly beyond the models solely on the coarse-grained end-to-end learning or geometry-based structural optimization alone.

## Results

The DRfold pipeline is outlined in **Figure 1**. The query sequence along with its SS predictions is first fed into an embedding layer which outputs the 1D sequence and 2D pair representations. The embedded representations then go through 48 RNA transformer blocks and are used for end-to-end RNA global-frame training, in terms of nucleotide-wise rotation matrices and translation vectors which can be used to recover the atomic coordinates of the RNA structure. Meanwhile, the pair representations are also used for RNA inter-nucleotide geometry prediction through a similar but independent set of transformer blocks. Finally, the predicted frame vectors and geometric restraints are aggregated into a composite potential for gradient-based RNA structure reconstruction, where the optimized conformation with the lowest energy is selected as the output model. DRfold was tested on 40 non-redundant RNA structures with length from 14 to 392 nucleotides, which are collected from sequence cluster centers (sequence identity cutoff of 90%) of solved structures deposited in or after the year 2021 in the PDB (Berman et al., 2000). Structures without any valid base pairs are not included. All the test sequences, as well as their cluster members, were excluded from the training dataset, which contains 3864 unique RNAs extracted from the PDB that were deposited before the year 2021. Thus, a filter based on both sequence identity and time stamp implements a stringent separation between the training and testing datasets of DRfold, both of which are available for download at https://zhanggroup.org/DRfold.

**Figure 1.**
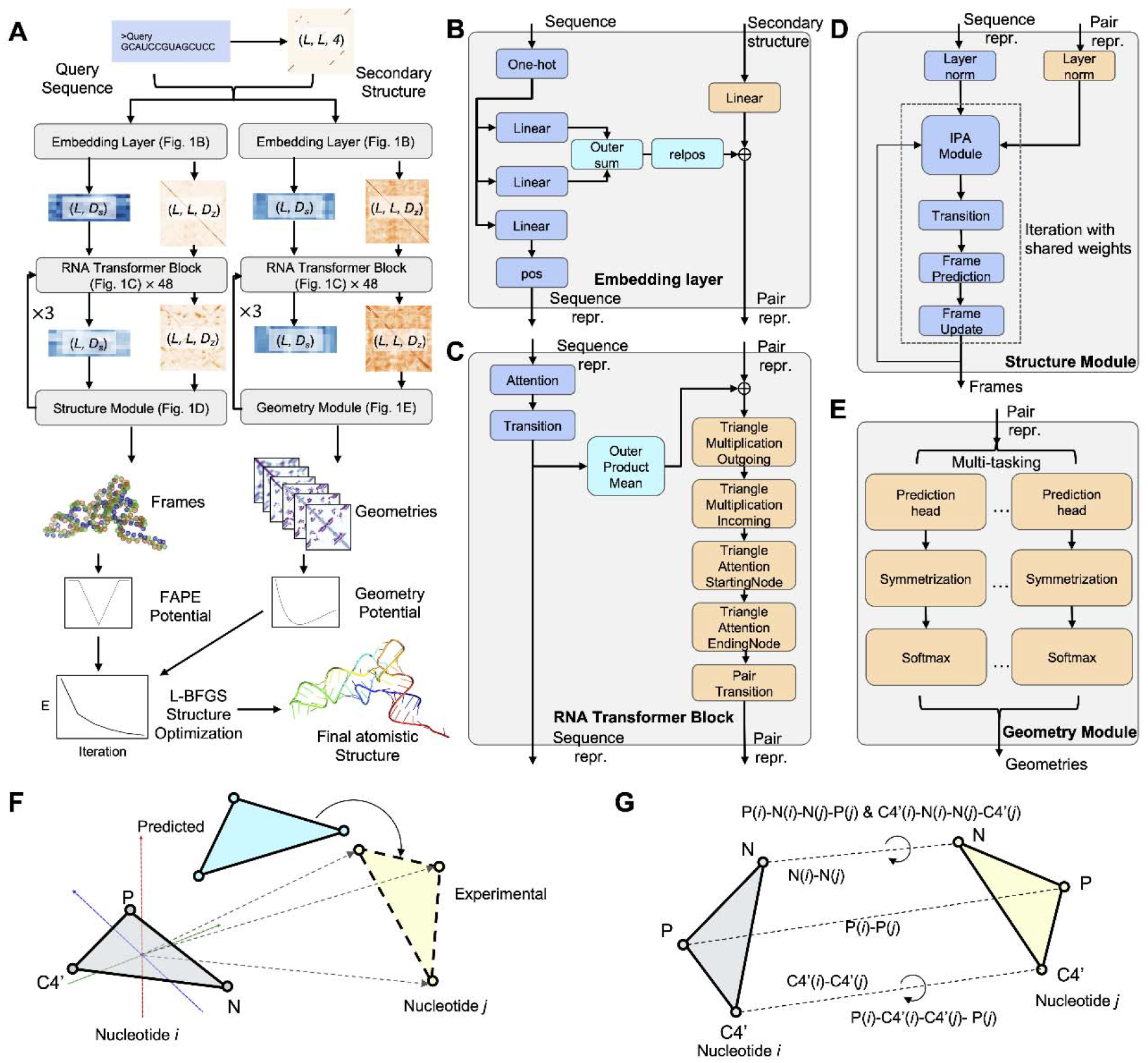
The pipeline of DRfold for deep learning-based RNA structure prediction by combining end-to-end model and geometry potentials. (A) DRfold pipeline for sequence-based RNA structure prediction. (B-E), Details of embedding layer, RNA transformer block, and structural and geometry modules respectively. (F) Illustration of the frame aligned point error (FAPE). (G) Prediction terms of inter-nucleotide geometry. and are hidden dimension sizes of sequence and pair features respectively, *L* is the length of the query sequence.

## DRfold outperforms current RNA structure predictors

To benchmark the performance of DRfold, two representative fragment assembly methods, RNAComposer (Biesiada et al., 2016) and 3dRNA (Zhao et al., 2012), and one representative *ab initio* RNA structure prediction method, RNA-BRiQ (Xiong et al., 2021), were considered as control methods. **Figure 2A** compares the root mean squared deviation (RMSD) of the models generated by DRfold and the control methods relative to the target structurewhere, where the coordinates of the P atoms are used for topology evaluation. The average RMSD value obtained by our method (12.76 Å) is significantly lower than those obtained by RNA-BRiQ (23.26 Å), RNAComposer (20.80 Å) and 3dRNA (20.54 Å), where the corresponding *P*-values are 7.34E-12, 1.02E-06, and 1.23E-07, respectively, in Student’s t-test. The median RMSD of DRfold is 6.89 Å, while the lowest median RMSD obtained in control methods is 19.23 Å (RNAComposor). Among the 40 test targets, 7 targets were found to have been successfully folded by DRfold at atomic resolution (RMSD < 2.5 Å). In **Figure 2B**, we further listed the accumulative fraction of cases with RMSD thresholds ranging from 2.5 Å to 15.0 Å, where DRfold generates significantly more cases than the control methods for all different RMSD cutoffs. For example, 55.0% of the DRfold models have a RMSD less than 7.5 Å, which is more than twice the fraction (20.0%) obtained by the best-performing third-party method, 3dRNA.

**Figure 2.**
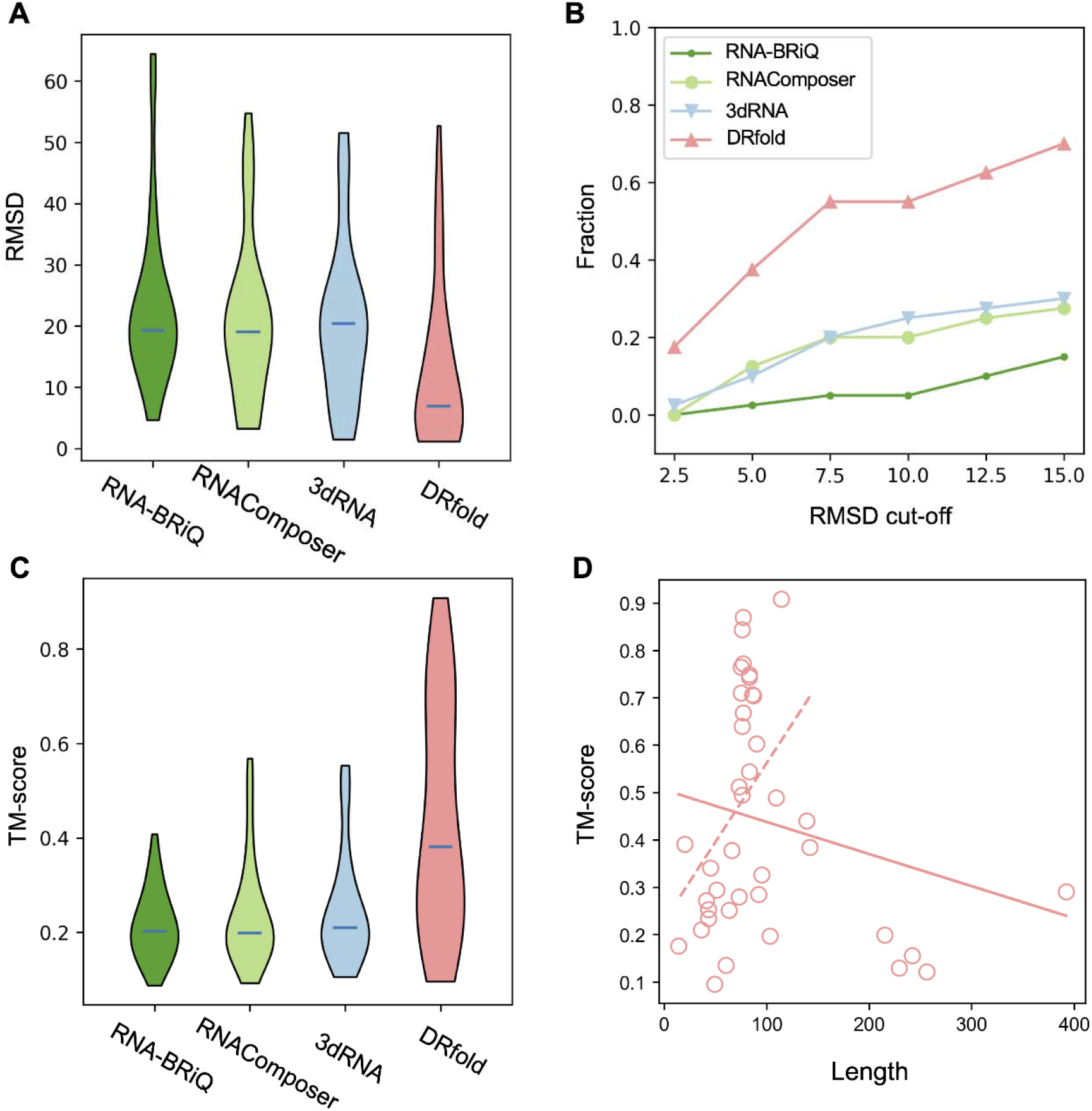
The comparison of DRfold with the control methods. (A) RMSD of the predicted models to the experimental structure. (B) Fraction of the test RNAs achieving successful structure prediction at different RMSD cut-offs. (C) TM-score distribution. (D) The correlation between the TM-score of DRfold models and the RNA length. Solid and dashed lines are fitted on all targets and the targets with length < 200 nucleotides, respectively.

Since a local error could cause a high RMSD, the RMSD value may not be ideal for assessing the quality of the RNA models at the high RMSD range. In **Figures 2C**, we further listed the results of TM-score, an index that is more sensitive to the global fold of the RNA models (Gong et al., 2019). Here, TM-score ranges in (0,1] with a higher value indicating a closer structural similarity, where a TM-score above 0.45 indicates a correct fold for RNA structures independent of the sequence length. As shown in **Figure 2C**, the average TM-score of the DRfold models (0.439) is 75.6% higher than the average TM-score of 0.250 obtained by the second-best method, 3dRNA, with a *P*-value of 6.74E-07. 42.5% (=17/40) of the RNA targets have the correct fold with TM-score > 0.45 predicted by DRfold, while the second-best method only achieves a success rate of 12.5%. The ability of DRfold to obtain very high-quality overall models for a substantial fraction of targets is apparent in the large upper shoulder in the distribution of TM-scores shown in **Figure 2C**. The advantage of DRfold is, however, not out of our expectation, as existing RNA structure prediction methods mostly utilize basic empirical and statistical potentials for *p* (*structure* |*sequence*). Given the limited number of parameters in their force fields, the global sequence conditions cannot be extensively counted and the generic potential forms (e.g., distance or angles) do not precisely determine the complex topology of the RNA structures. In contrast, the extensive weighting parameter setting embedded in the transformer module used by DRfold allows for access to the global sequence information. In addition, the end-to-end loss function (see Methods) can further ensure the high correspondence of deep learning prediction and the correct overall conformation. In **Figure 2D**, we listed the scattering plot of TM-score versus the length of the test RNAs, where a weak correlation (Pearson Correlation Coefficient, PCC = −0.20) can be observed, indicating that the performance is overall weakly dependent on the RNA length. It is notable that for those targets with length > 200 NTs, the predictions of DRfold are not satisfying. One possible reason is that a maximum RNA length has been set to 200 when we trained the models in DRfold, suggesting that the interaction patterns for extreme long-distant (>200) nucleotide pairs may not be sufficiently learned. Interestingly, after excluding the 5 extreme targets with length > 200, the PCC becomes extraordinarily different, i.e., 0.71 (see dashed line in **Figure 2D**). Such correlation would suggest that the performance will be better with longer (< 200) RNA targets. This is probably due to the design of the loss function used in training the end-to-end models (see Eq. 2 in Methods), where the summation of the FAPE loss for all nucleotide pairs puts more emphases on larger RNAs with the weight factor of *L*^2^ multiplied to the mean FAPE loss, compared to the weight factor of 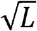 in AlphaFold2 (Jumper et al., 2021). In fact, RNA structures with larger size do naturally contains more information. It has been observed in our validations that using the summed loss function results in a much better performance than using the average of FAPE loss over the pairs, while training with mean FAPE loss may fail to converge.

In **Figure 3**, we presented a detailed head-to-head comparison of RMSD and TM-score between DRfold and the control methods, where a pronounced advantage of DRfold with the control methods was observed in all the boxes. For example, the fraction of the test targets for which DRfold achieves a lower RMSD than the control methods are 95.0% (to BRiQ), 82.5% (RNAComposor) and 90.0% (3dRNA), respectively, as shown in **Figures 3A-C**. The superiority of DRfold is more robust when evaluated by TM-score in **Figures 3D-F**, as the maximum absolute difference in TM-score is only 0.036 for those targets where any of the control methods perform better. In contrast, for those targets where DRfold have a higher TM-score, the absolute difference can be up to 0.742. Such observation suggests again that DRfold can consistently predict better global conformations compared to the classic RNA folding methods.

**Figure 3.**
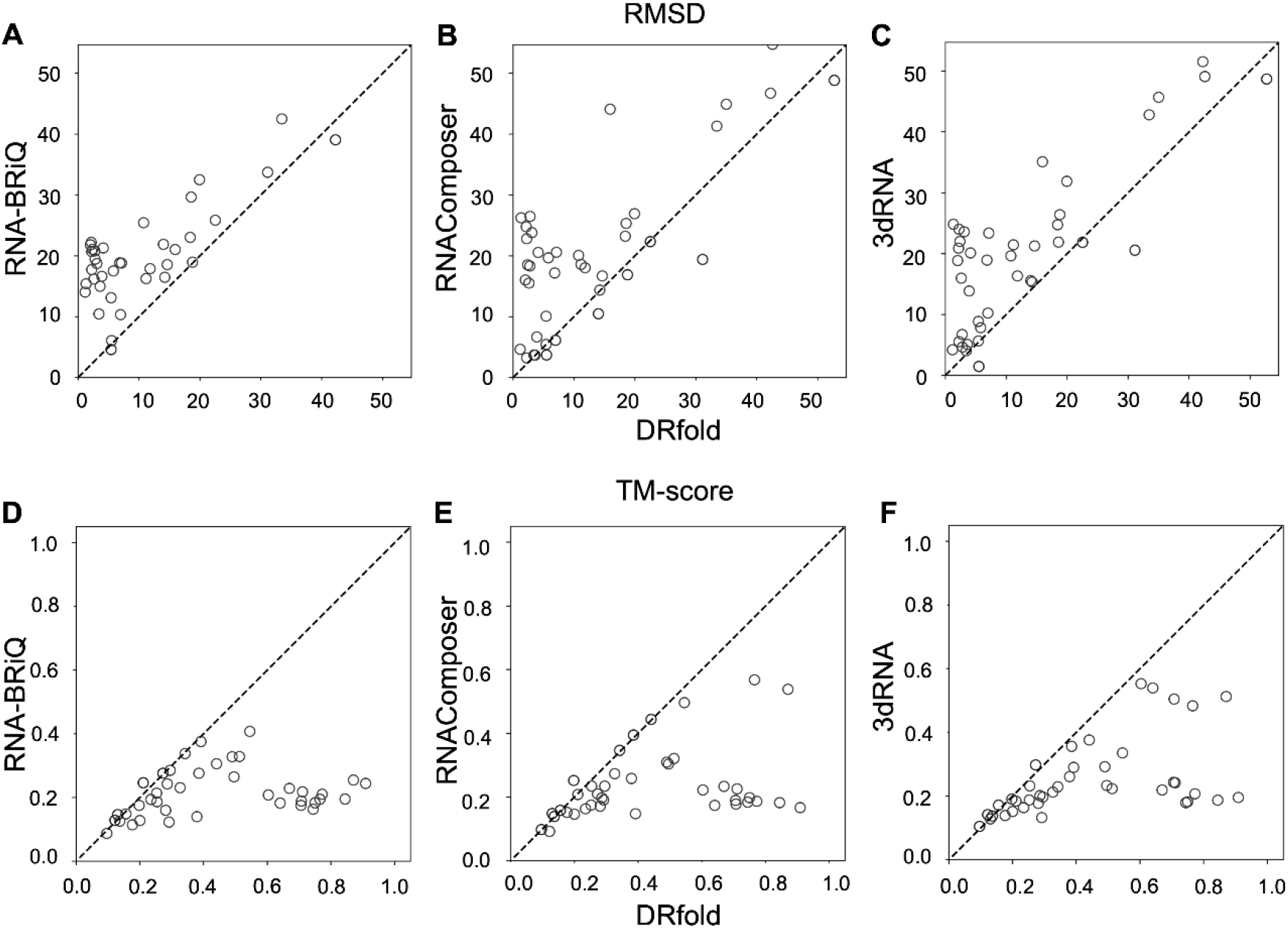
Head-to-head comparisons between DRfold and the control methods. (A-C) Head-to-head comparisons of RMSDs. (D-E) Head-to-head comparisons of the TM-scores.

### End-to-end models provide complementary information to geometric restraints for RNA structure modeling

The core of the DRfold pipeline is the introduction of two types of complementary potentials, i.e., FAPE potential and geometry potentials, from two separate transformer networks. The former models directly predict the rotation matrix and the translation vector for the frames representing each nucleotide, forming an end-to-end learning strategy for RNA structure. In DRfold, 6 independent end-to-end models were trained with different parameter initializations. **Table S1** lists the average TM-score of those models on the test set. Without any post-processing, the individual end-to-end models already outperform all control methods significantly. For example, the lowest average TM-score (0.393) obtained among 6 end-to-end models is 57.2% higher than that of the best control method 3dRNA (0.250). After applying an optimization procedure which is an ensemble of the 6 conformations, the average TM-score rises to 0.417.

To further examine the importance of the end-to-end potential to DRfold, we plot in **Figure 4A** a TM-score comparison of the models by the full-version DRfold with that removing the FAPE potential in DRfold, where the latter means that the structures are only optimized by the geometry potentials. On average, the TM-score drops from 0.439 to 0.413, with a *P*-value of 2.7E-02 in Student’s t-test, indicating that the performance loss is statistically significant. In **Figure 4C-E** we present one example from the sgRNA (PDB ID: 7OX9 Chain A) in the Cas9 endonuclease. The model built on the geometry potential has a reasonable fold but with significant local errors mainly in the regions of 5’- and 3’-terminal regions and the central loop (26-41 NTs), which resulted in an overall TM-score=0.369 and RMSD=6.52 Å. As shown in **Figure 4E**, the end-to-end structural models have variable quality in the 5’- and 3’-terminal regions and with consistently lower error in the loop region. A consensus-based optimization of both end-to-end and geometric potentials resulted in a significantly improved model with a TM-score=0.749 and RMSD=2.00 Å (see **Figure 4D** and the bottom of **Figure 4E**).

**Figure 4.**
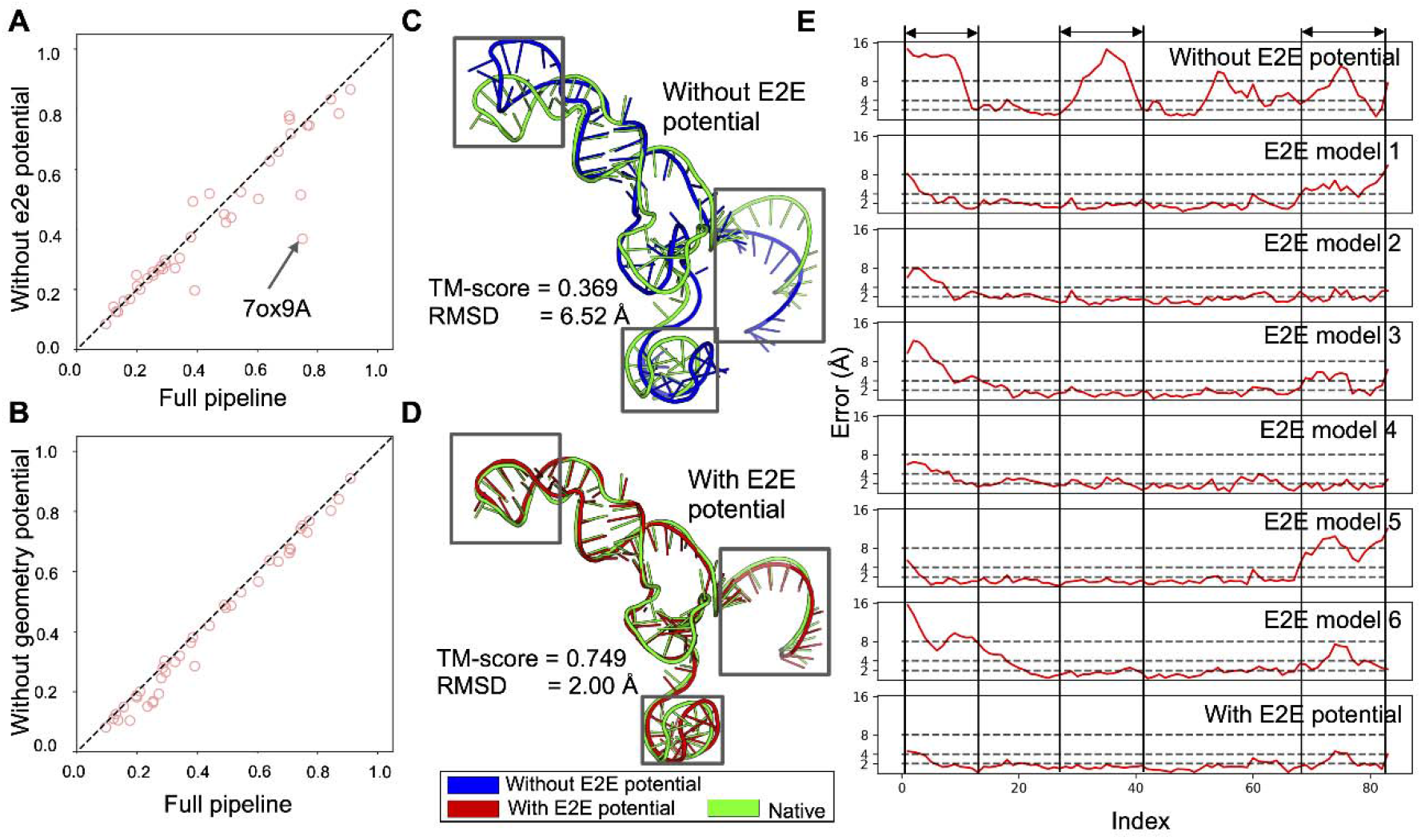
Performance comparisons between the full DRfold pipeline and those without component potentials. Comparison of full model performances vs. reduced models (A) without end-to-end potential and (B) without geometry potential. (C) Structure superposition between experimental structure (green) and the structure predicted without end-to-end potential (blue) for target 7OX9A. (D) Structure superposition between experimental structure (green) and the structure with the full DRfold pipeline (red) for target 7OX9A. (E) Residue-wise errors of the structure predicted without end-to-end potential, the structures predicted by 6 end-to-end models, and the structure predicted by the full DRfold pipeline for target 7OX9A. The residue-wise errors were computed from the superpositions produced by TM-score.

Here, the geometry potential adopts a composite set of terms representing inter-nucleotide geometry, including distances and torsion angles. To examine the impact of such potential to DRfold, we compare the TM-scores of DRfold on the 40 test RNAs with and without geometry potentials in **Figure 4B**. It was observed that including the geometry potentials on top of end-to-end potential brings small but consistent improvements in TM-score (with *P*-value=2.1E-07). In **Figure S1**, we listed the TM-score improvements by the geometry potentials over the RNA length, which shows that the incorporation of the additional long-range inter-nucleotide geometric restraint potential can consistently improve the folding performance for RNA with various sizes.

Overall, these results demonstrate that, although they start from the same set of sequence and SS embedding matrices in the network, the independent training of the end-to-end and geometric potentials have learned structural features complementary to each other and collectively improve the overall quality of the DRfold models.

### Secondary structure prediction facilitates feature learning and model construction

Unlike the strategy used in AlphaFold2 (Jumper *et al*., 2021) which feds the embedding module directly with the query sequence, DRfold uses a consensus of two SS predictors by RNAfold (Lorenz et al., 2011) and PETfold (Seemann et al., 2008) to extract 2D features for the additional pair embedding. To examine the impact of the SS predictions, **Figure 5A** shows a head-to-head comparison of TM-score for the full DRfold pipeline vs. an ablated pipeline in which the predicted SS input is omitted. There is a significant performance drop, i.e., from 0.439 to 0.295 in TM-score, without secondary structure.

**Figure 5.**
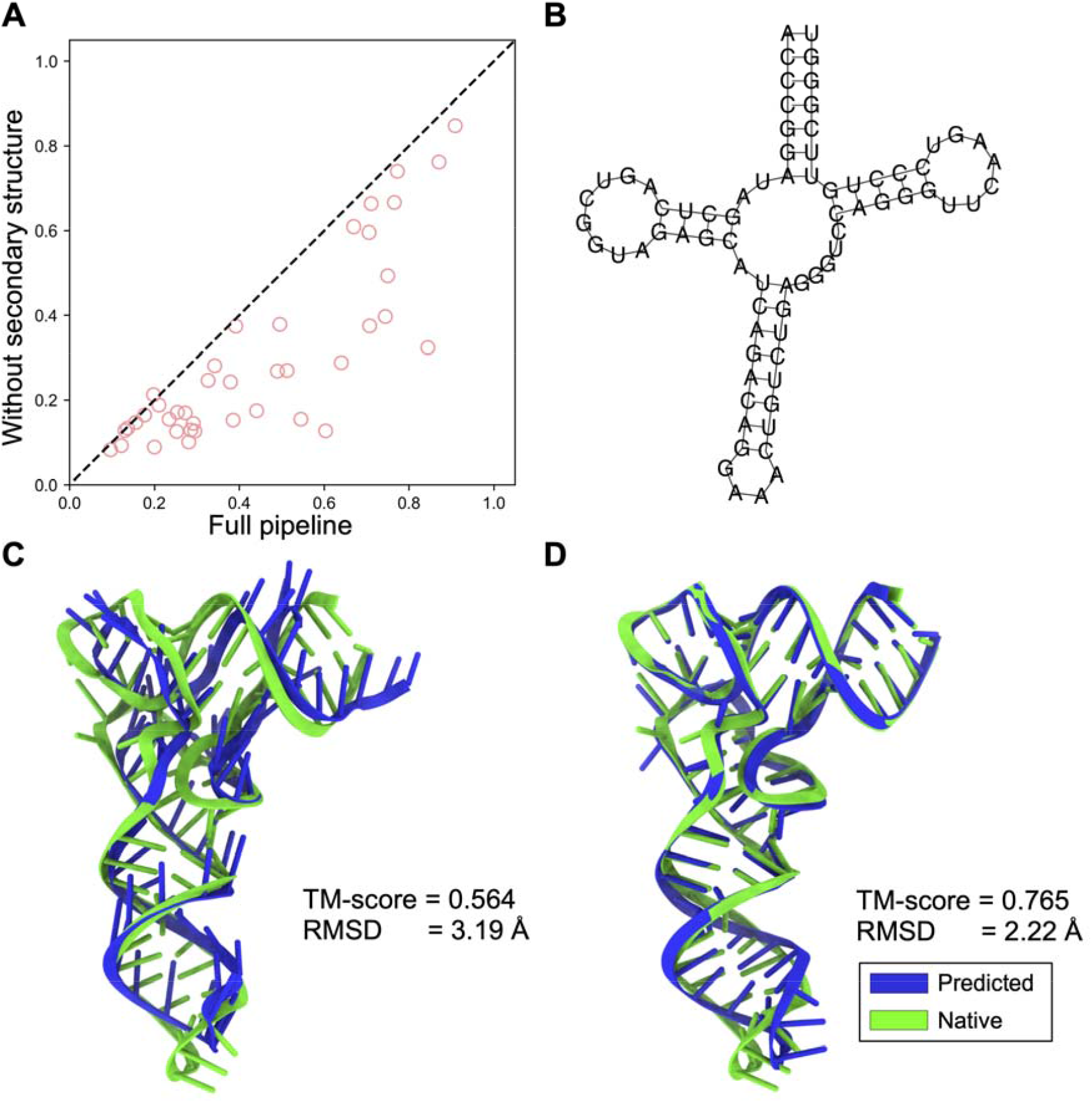
Secondary structure feature improves performance. (A) TM-score comparisons between the full DRfold pipeline and a reduced model without predicted secondary structure information. (B) The predicted secondary structure for target 7MRLA. (C) Superposition of the predicted structure (blue) by DRfold using only the secondary structure with the experimental structure (green) for target 7MRLA. (D) Superposition of the predicted structure (blue) by the full version DRfold with the experimental structure (green) for target 7MRLA.

As a complementary test to determine the amount of information on tertiary structure that can be obtained from secondary structure alone, we performed an additional experiment in which we replaced all input residues with an “N” (which represents the “unknown” residue type) and only fed DRfold with predicted secondary structure features, mimicking a tertiary structure recovery task from SS prediction only. Unsurprisingly, the average TM-score significantly drops from 0.439 to 0.268. Nevertheless, the TM-score value is still considerably higher than that by the best third-party program 3dRNA based on statistical models (0.250). Among the 40 test targets, there are 5 targets that have a correct fold with TM-score > 0.45 on the sequence-free DRfold modeling.

In **Figure 5B-C**, we show one example in which DRfold successfully recovered the overall topology only based on its predicted secondary structures. This is a tRNA (PDB ID: 7MRL Chain A), which has a clover-like structure. Based on the predicted SS only (**Figure 5B**), DRfold constructs a model of correct fold with TM-score=0.564 and RMSD=3.19 Å (**Figure 5C**). Nevertheless, we noticed that many of the base-pairs are not correctly formed, probably due to the lack of nucleotide sequence information. After the inclusion of the sequence information, the RNA model by the full DRfold pipeline has a much-improved base-pairing quality with the overall TM-score=0.765 and RMSD=2.22 Å (**Figure 5D**), demonstrating the impact of sequence-specific base-pairing on the RNA structure modeling.

Overall, these results demonstrated the significant importance of the SS embedding feature to the DRfold performance, although the pipeline requests the nucleotide sequence only. In principle, an ideal deep learning model should be able to learn the secondary structure directly from sequence. Previous studies (Chen et al., 2020; Singh et al., 2019) have showed the success in learning the RNA secondary structures from sequences. However, given the limitation of available RNA structure data, relevant input structural feature, containing auxiliary information related to the RNA topology such as SS, should be greatly beneficial to facilitate the neural networks to improve the learning efficiency and effectiveness in RNA tertiary structure prediction.

### Blind RNA structure prediction in CASP15

An automated program of DRfold participated in the most recent community-wide CASP15 experiment for RNA structure prediction with a Group ID of ‘rDP’. Although there were only 12 test targets (Das, 2022), this gives an opportunity to objectively assess DRfold relative to the state of the art of the field. In **Table S2 and S3**, we list the cumulative Z-scores for all groups in terms of RMSD and TM-score respectively. Following the convention of CASPs, the Z-scores were calculated in the following procedure: (1) for given target raw Z-scores are calculated as the difference between the raw score and the mean in the unit of standard derivation for the first models of different groups; (2) remove the outlier models with raw Z-scores below the tolerance threshold (set to −2.0); (3) recalculate Z-scores on the reduced model set; (4) assign Z-scores below the penalty threshold (either −2.0 or 0.0) to the value of this threshold. As shown in **Table S2**, using RMSD Z-score (calculated by negative of RMSD values), DRfold ranks 5^th^ and 6^th^ with penalty thresholds of −2.0 and 0.0 respectively. When using TM-score, the ranking becomes 6^th^ and 9^th^ respectively (**Table S3**).

There is an obvious performance gap between DRfold and the top 4 methods (AIchemy_RNA2, Chen, RNApolis and GeneSilico), which fold RNAs guided by highly specialized human-expertise in the field of RNA structure. Our method, in contrast, only requires single sequence information and is fully automatic. Nevertheless, we find that the performance of DRfold is comparable with that of other top methods which requires additional information sources such as templates (e.g., CoMMiT-server), MSAs (e.g., AIchemy_RNA, Yang-Server and UltraFold) or pretrained nucleotide sequence models (e.g., AIchemy_RNA), especially when evaluating with RMSD. Considering that the only input of DRfold is the RNA sequence, the reason for such competitive performance should be attributed to the network of DRfold that learns complementary potentials to improve RNA folding. Meanwhile, the excellent performance of other top-ranked methods demonstrated potentials to further improve DRfold with the integration of those additional information sources.

## Discussion

We developed a novel method, DRfold, for *ab initio* RNA structure prediction based on single sequence-based deep learning models. The approach is able to learn the coarse-grained RNA structures directly from sequence in an end-to-end fashion, using cutting-edge self-attention transformer networks. The predicted conformations are further optimized by a separately trained deep-geometric potential through gradient-descent based simulations. The method was tested on a nonredundant set of RNA structures, which are separated from the training RNAs with a strict control of release time and sequence identity cutoff, where the results showed significant advantage over existing approaches built on statistical model and fragment assembly simulations.

The success of DRfold mainly arises from the deep learning-based potentials, which, to our knowledge, have rarely been introduced in existing RNA structure prediction pipelines. The end-to-end models in DRfold have proven to be highly effective in predicting the frames of residue positions that can recover the atomic model of RNA structures through trained rotation and translation matrices. With the integration of geometry restraints, the hybrid potentials can further improve the accuracy for structural models through atomic-level optimization. Moreover, the predicted secondary structure features from physical based folding programs were found beneficial to facilitate the network learning and help generate more accurate base-pairing and local structural packing for the RNA models.

Despite the success demonstrated here, we find that the overall performance of structure prediction for RNA is still limited, compared to that for proteins (for example, AlphaFold2 (Jumper et al., 2021)). This may be partly due to the lack of sufficient number of RNA structures that are needed to train the high number of parameters in the end-to-end networks. This is especially true for the CASP15 targets (Das, 2022), where DRfold performed better on the nature RNAs than the synthetic RNAs that may bear different folding pattern from the nature RNAs on which DRfold was trained. Meanwhile, to facilitate the modeling of hard RNAs without many homologous sequences, DRfold has been trained only on single sequence, while the inclusion of multiple sequence alignments (Zhang et al., 2020) might also help to improve the RNA structure prediction through the aggregation of more extensive evolutionary features. Nevertheless, the release of the open-source DRfold program provides a useful platform to the community for efficient deep learning-based RNA structure prediction whose accuracy will continue to improve with the progress of new machine learning techniques and RNA structure and sequence databases.

## MATERIALS AND METHODS

DRfold is a deep machine learning based approach to *ab initio* RNA structure prediction. It consists of two steps of end-to-end frame and geometric potential training, and deep-potential guided full-atom model construction, where the general pipeline of the method is shown in **Figure 1**.

### Feature preparation and embedding

The only required input of DRfold is the nucleotide sequence, which is represented by a 5-D one-hot encoding, including 4 types of nucleotides (‘A’, ‘U’, ‘G’, ‘C’) and an unknown state (‘N’) that can represent modified or degenerate nucleotides. Based on the sequence, the SS is predicted by two complementary methods: RNAfold (Lorenz et al., 2011) and PETfold (Seemann et al., 2008), which are concatenated in the network. The predicted SS is in the form of the matrix where the entry is set to 1 if the corresponding residue pair forms a base pair. We also include the SS probability map predicted by the considered methods, which brings in another 2 channels for pair input. Thus, given a sequence of length *L*, the sequence feature (1-D) and the pairwise feature (2-D) have the shape of *L* × 5 and *L* × *L* × 4, respectively, where ‘4’ is from 4 SS channels (**Figure 1A)**.

The sequence feature *s*_*F*_ ∈ *R*^*L*×5^ and the pair feature *Z*_*F*_ ∈ *R*^*L*×*L*×4^ act as the input of the embedding layer. The sequence feature *S*_*F*_ will be projected to the desired dimension (D_*S*_=64) by a linear layer. Another two linear projections of *S*_*F*_ will be added vertically and horizontally to form the initial pair representation. The initial pair representation will then be added to the projected representation of pair feature *Z*_*F*_, with a channel size of D_*Z*_ = 64. Thus, the output of the embedding layer contains the sequence representation *S* ∈ *R*^*L*×64^ and the pair representation *z* ∈ *R*^*L*×*L*×64^ **Figure 1B**)

We also embedded the 1-D and 2-D positional encodings (‘pos’ and ‘relpos’ layers in Figure 1B) to *s* and *z*. A recycling strategy is used by encoding the geometry descriptors of the predicted structure conformation (for end-to-end models only), bringing the 1-D representation and the 2-D representation from the previous recycle to the input of the current recycle. The recycle number is set to 3 for both end-to-end and geometry models (**Figure 1A**).

### RNA transformer network

There are a total of 48 transformer blocks in DRfold. The transformer block module is extended from the design of Evoformer in AlphaFold2 (Jumper et al., 2021). As shown in **Figure 1C**, the sequence representation *S* and the pair representation *Z* will first go through a sequence row-wise gated self-attention with pair bias module that output the new sequence representation. The number of heads and the channel size of each head are set to 8 and 8 respectively. A sequence transition layer that contains 2 linear layers will be stacked after the sequence self-attention. The sequence transition layer first expands the dimension from 64 to 128 and then projects it to the original channel size (64). The obtained sequence representation is transformed to a pair representation by an outer product mean (OPM) block. The OPM block first projects the sequence channel size to 12 with two separated linear layers. After the outer product mean operation, the output 2-D representation thus has a channel size of 12 × 12. Another linear layer in the OPM block then projects it to the desired channel size, i.e., 64.

The 2-D representation from the OPM block will sequentially go through a set of blocks, including (1) Triangle multiplicative update block using outgoing edges, (2) Triangle multiplicative update block using incoming edges, (3) Triangle self-attention block around starting node, (4) Triangle self-attention block around starting node, and (5) pair transition block. For Triangle multiplicative update blocks, the channel size is set to 32; for Triangle self-attention blocks, the number of head and the channel size are set to 4 and 8, respectively. It should be noted that all the sequence and pair blocks are stacked residually (He et al., 2016) for efficient and stable training.

### RNA structure module

The end-to-end model in DRfold predicts the spatial location of each nucleotide, which can be represented by a rotation matrix and a translation vector operating on a predefined conformation at a local frame. Here, we used three atoms (i.e., P, C4’, and the glycosidic N atom of the nucleobase) to represent the coarse-grained conformation of a nucleotide. We assume that the three atoms form a rigid body for each of the four nucleotides. The RNA structure module takes the sequence representation from the RNA transformer network as the input to iteratively train the nucleotide-wise rotation matrices and translation vectors. As shown in **Figure 1D**, the pair representation is also utilized by the invariant point attention (IPA) module to equivariantly update the RNA structure conformation in the iterations. The channel sizes of sequence and pair representations are set to 128 and 64, respectively, in the RNA structure module. The hyper-parameters in IPA, (*N*_*head*_,*c,N*_*querypoints*_,*N*_*pointvalues*_), are set to (8, 16, 4, 6), and the iteration number is set to 5.

An important step of the RNA structure module is the construction of the local frames from ground truth positions. Considering the higher flexibility of RNA structures compared to that of proteins, we construct frames with the SVD orthogonalization (Aiken et al., 1980), instead of the Gram–Schmidt orthogonalization that was used in Alphafold2 (**Figure S2**). Since SVD orthogonalization maximizes the likelihood in the presence of Gaussian noise, it is less greedy than the Gram–Schmidt orthogonalization and thus has a lower bias during the estimation process. Levinson et al. (Levinson et al., 2020) has showed that in the view of matrix reconstruction, the approximation errors of SVD orthogonalization are half that of the Gram-Schmidt procedure.

### Loss function of end-to-end models

Two types of loss functions, including the frame aligned point error (FAPE) loss (**Figure 1F**) and the inter-N atom distance loss, are used when training the end-to-end models, i.e.,

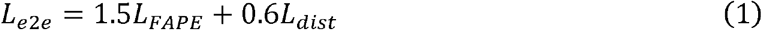

The FAPE loss is defined by

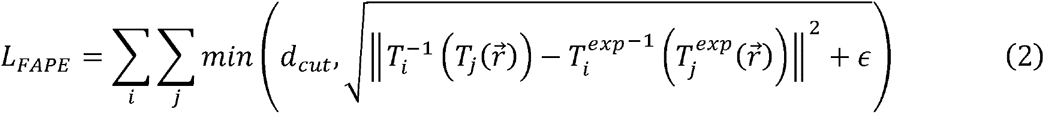

where *T* (or 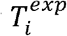) represents the Euclidean transformation, including rotation matrix and translation vector that will be learned in the networks, to convert a position at the local frame 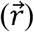 to the global space, for the predicted model (or the target experimental structure in the training dataset). The parameter *d*_*cut*_ is set to 30 Å and *ϵ* is set to 10 ^− 3^ Å. Here, *I* and *j* enumerate all the nucleotide positions and all the atoms of the RNA structure, respectively. Because each term contains two reversible transforms, i.e 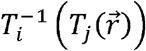, any rigid-body transformations will be cancelled out in the calculation. Therefore, *L*_*FAPE*_ is by design invariant to any rigid-body conformational transformations to the predicted structures.

The distance loss function *L*_*dist*_ in Eq. (1) takes the cross-entropy form of

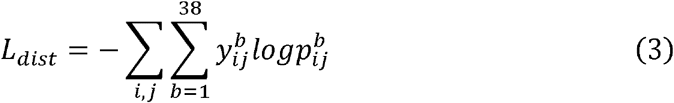

where 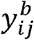 is an indicator function to check if the distance of atom pair (*i,j*) of the target experimental structure falls into *b*-th distance interval; and 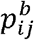 is the predicted probability for the interval. The inter-atom distance is split into 36 intervals between 2-40 Å, with additional two bins representing distance < 2 Å and >40 Å.

### Prediction terms of geometry models

For a pair of residues, a set of geometry potentials are extracted from the experimentally determined structures as supervised information to train deep geometric potentials (Li et al., 2022). As shown in **Figure 1G**, the Euclidean distance between P, C4’ and N atoms are calculated, where the distance values of inter-P atoms, inter-C4’ atoms and inter-N atoms are discretized into 56, 44, and 32 bins in the ranges of [2, 30 Å], [2, 24 Å] and [2, 18 Å], respectively. For each distance term, two addition bins are added representing values < 2 Å and >*M* Å, where *M* is the corresponding maximum distance values (30, 24, and 18 Å, respectively). Meanwhile, the long-range dihedral angles formed by atoms of each nucleotide pair (*i, j*) will be extracted, which are formed, respectively, by P(*i*)-C4’(*i*)-C4’(*j*)-P(*j*), C4’(*i*)-N(*i*)-N(*j*)-C4’(*j*) and P(*i*)-N(*i*)-N(*j*)-P(*j*). The dihedral angle values are discretized into 36 bins, plus the dimension representing whether the length of virtual bond, i.e., C4’(*i*)-C4’(*j*) and N(*i*)-N(*j*), is larger than their corresponding maximum distance values *M* (=24 and 18 Å, respectively).

The loss function of geometry models is the cross-entropy loss of distance and dihedral angle terms defined by

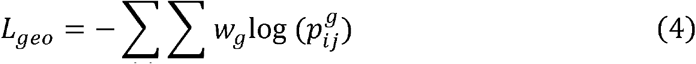

where *G* is the set of geometry terms of the distance and dihedral angles, and weight parameters *w*_*g*_=1.0 and 0.5 for distance- and dihedral angle-related losses, respectively. The models 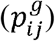 are trained at a multi-task learning architecture as described in **Figure 1E**.

### Training of the end-to-end and geometry models

The end-to-end and geometric network models were trained by the Adam optimizer (Kingma and Ba, 2014) with an initial learning rate of 1e-3 for 100 epochs. The maximum RNA sequence length was set to 200. For an RNA sequence with over 200 nucleotides, a continuous segment of 200 nucleotides was randomly sampled during the training. The batch size was set to 3 to accelerate the training with the gradient accumulation mechanism in PyTorch (Paszke et al., 2019). We also use gradient checkpointing to reduce the memory occupancy for each transformer block (Chen et al., 2016). The whole end-to-end model was trained on a single Nvidia A40 GPU with 32GB of memory, where 6 end-to-end models and 3 geometry models with different random parameter initializations were trained, and training each of them took 2 weeks. For the 3 geometry models, it took around 50 epochs of training for 5 days for each.

### Structure optimization with integrated end-to-end and geometry potentials

Following the end-to-end and geometry modeling, a combination of two deep-learning energy terms, *E*_*DL*_ = *E*_*e*2*e*_ + *E*_*geo*_, is used to guide the next step of RNA structure optimization. The first end-to-end potential is written as

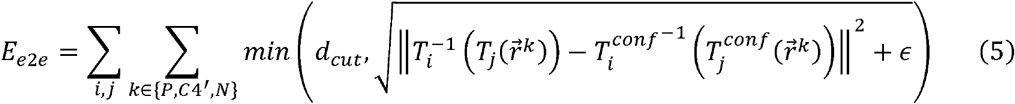

where *T*_*i*_ is the predicted rotation matrices and translation vectors by the end-to-end models as defined in Eq. (2) and are kept unchanged during the structure optimization. 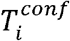 represent the transforms to recover the RNA conformation of the predicted model for the atom set {*P,C*4’,*N*} from the predefined local frame, 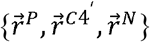. We sum the end-to-end energy values calculated by 6 end-to-end models as the final consensus end-to-end potential.

The geometry potential *E*_*geo*_ is defined by

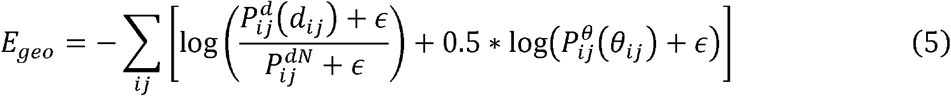

where 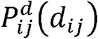 and 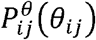 are the predicted probabilities for a given distance *d*_*ij*_ and dihedral angles θ_*ij*_ between a nucleotide pair 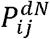 and 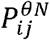 are corresponding probability of the last distance bin below the upper threshold. The negative log-likelihood of the predicted probabilities is interpolated by cubic spline to form a potential curve for the specific distance/dihedral angle terms (Li et al., 2022).

The 6 conformations predicted by end-to-end models were also used as initial structures of the optimization system and separately optimized by the same hybrid potential function. The gradient of parameters with respect to the hybrid potential function can be calculated by the automatic differentiation package in PyTorch. With the energy value and the gradient, we can use the L-BFGS algorithm (Zhu et al., 1997) to iteratively update the parameters of the system, i.e., 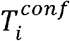, which determines the 3D conformations of the RNA models. The conformation with lowest energy is considered as the final predicted structure, among the 6 different initials.

Since both the end-to-end pipeline and the L-BFGS simulation operate only on the P, C4’ and N atoms, we need to reconstruct the full atomic model from the 3-atom coarse-grained model. For this, the standard conformations of the full-atomic structure for each of the four types of nucleotides are extracted from the generic A-form RNA helix with a 32.7° twist and a 2.548 Å rise (Chandrasekaran and Arnott, 1989). The standard conformation of the full-atomic nucleotide is then superimposed to the three-atom model by the least square fit to quickly obtain the final full-atomic RNA structure.

## ACKNOWLEDGMENTS

We thank Drs. Sha Gong and Xi Zhang for insightful discussions. This work was supported in part by the National Institute of General Medical Sciences (GM083107, GM116960, GM136422 to Y.Z.); the National Institute of Allergy and Infectious Diseases (AI134678 to P.L.F. and Y.Z.); the National Institute of Health Office of The Director (OD026825 to Y.Z.); the National Science Foundation (DBI2030790, IIS1901191, MTM2025426 to Y.Z.). This work used the Extreme Science and Engineering Discovery Environment (XSEDE), which is supported by National Science Foundation (ACI1548562).

## Author contributions

Y.Z. conceived and designed the project and supervised the work. Y.L. developed the methods and performed the benchmark. C.Z. prepared the data and developed the full atom packing software. C.F. prepared the initial data. P.L.F. supervised the work. All authors wrote the manuscript and approved the final version.

## Data and materials availability

All data needed to evaluate the conclusions in the paper are present in the paper and/or the Supplementary Materials. The DRfold server is available at https://zhanggroup.org/DRfold/.

