## Supplementary Material for "Integrating end-to-end learning with deep geometrical potentials for *ab initio* RNA structure prediction"

#### Supplementary Tables

**Table S1. Benchmark results of 6 end-to-end models and their ensemble.** The *P*-values are computed according to Student's t-test.

| Models | Mean TM-score | Median TM-score | *P*-value |
|---|---|---|---|
| Model 1 | 0.395 | 0.335 | 2.4e-02 |
| Model 2 | 0.398 | 0.331 | 6.6e-02 |
| Model 3 | 0.393 | 0.320 | 1.4e-02 |
| Model 4 | 0.394 | 0.346 | 4.5e-02 |
| Model 5 | 0.405 | 0.332 | 1.8e-01 |
| Model 6 | 0.397 | 0.342 | 4.0e-02 |
| Ensemble | **0.417** | **0.372** | - |

**Table S2.** Z-score based relative group performance of first model for RMSD with penalty thresholds of -2.0 and 0.0 respectively.

| SUM Z-score > -2.0 | | | SUM Z-score > -0.0 | | |
|---|---|---|---|---|---|
| Rank | Group ID | SUM Zscore | Rank | Group ID | SUM Zscore |
| 1 | Chen | 13.46 | 1 | Chen | 15.00 |
| 2 | AIchemy_RNA2 | 13.40 | 2 | AIchemy_RNA2 | 14.48 |
| 3 | RNApolis | 10.74 | 3 | RNApolis | 11.22 |
| 4 | Yang-Server | 06.01 | 4 | GeneSilico | 08.14 |
| **5** | **rDP** | **05.72** | 5 | Yang-Server | 06.68 |
| 6 | CoMMiT-server | 03.63 | **6** | **rDP** | **06.18** |
| 7 | CoMMiT-human | 03.48 | 7 | AIchemy_RNA | 05.73 |
| 8 | UltraFold | 02.14 | 8 | UltraFold | 05.73 |
| 9 | Yang | 01.97 | 9 | Yang- Multimer | 05.27 |
| 10 | Kiharalab | 01.50 | 10 | CoMMiT-server | 05.21 |
| 11 | UltraFold_Server | 00.68 | 11 | CoMMiT-human | 05.11 |
| 12 | GeneSilico | 00.56 | 12 | Yang | 04.83 |
| 13 | AIchemy_RNA | 00.38 | 13 | Kiharalab | 04.69 |
| 14 | Yang- Multimer | -00.39 | 14 | UltraFold_Server | 04.31 |
| 15 | Coqualia | -02.41 | 15 | SoutheRNA | 03.42 |
| 16 | SoutheRNA | -02.68 | 16 | LCBio | 03.29 |
| 17 | LCBio | -02.90 | 17 | Coqualia | 03.24 |
| 18 | BAKER | -04.08 | 18 | DF_RNA | 02.67 |
| 19 | Rookie | -04.49 | 19 | BAKER | 02.64 |
| 20 | Manifold-E | -06.69 | 20 | nucE2E | 02.62 |
| 21 | SHT | -06.78 | 21 | Rookie | 01.91 |
| 22 | GinobiFold | -06.97 | 22 | CoDock | 01.75 |
| 23 | FoldEver | -07.49 | 23 | AIchemy_LIG | 01.74 |
| 24 | GWxraylab | -07.91 | 23 | AIchemy_LIG3 | 01.74 |
| 25 | FoldEver-Hybrid | -08.61 | 23 | AIchemy_LIG2 | 01.74 |
| 26 | Manifold | -09.04 | 26 | PerezLab_Gators | 01.53 |
| 27 | DF_RNA | -10.58 | 27 | Manifold | 01.37 |
| 28 | nucE2E | -11.38 | 28 | SHT | 01.23 |
| 29 | CoDock | -12.25 | 29 | FoldEver | 01.20 |
| 30 | Schug_Lab | -12.90 | 29 | FoldEver-Hybrid | 01.20 |
| 31 | PerezLab_Gators | -16.38 | 31 | GinobiFold | 01.11 |
| 32 | WL_team | -19.33 | 32 | Venclovas | 01.00 |
| 33 | Graphen_Medical | -19.37 | 33 | WL_team | 00.86 |
| 34 | Kiharalab_Server | -19.48 | 34 | Manifold-E | 00.66 |
| 35 | Venclovas | -19.48 | 35 | Schug_Lab | 00.55 |
| 36 | AIchemy_LIG | -20.26 | 36 | Kiharalab_Server | 00.49 |
| 36 | AIchemy_LIG3 | -20.26 | 37 | GWxraylab | 00.49 |
| 36 | AIchemy_LIG2 | -20.26 | 38 | Manifold-LC-E | 00.33 |
| 39 | Manifold-LC-E | -21.67 | 39 | UNRES | 00.00 |
| 40 | Manifold-LC | -22.53 | 39 | Manifold-LC | 00.00 |
| 41 | UNRES | -24.00 | 39 | Graphen_Medical | 00.00 |

**Table S3.** Z-score based relative group performance of first model for TM-score with penalty thresholds of -2.0 and 0.0 respectively.

| SUM Z-score > -2.0 | | | SUM Z-score > -0.0 | | |
|---|---|---|---|---|---|
| Rank | Group ID | SUM Zscore | Rank | Group ID | SUM Zscore |
| 1 | AIchemy_RNA2 | 20.72 | 1 | AIchemy_RNA2 | 21.35 |
| 2 | Chen | 16.34 | 2 | Chen | 16.42 |
| 3 | RNApolis | 12.44 | 3 | RNApolis | 12.91 |
| 4 | GeneSilico | 04.28 | 4 | GeneSilico | 10.48 |
| 5 | Yang-Server | 02.83 | 5 | AIchemy_RNA | 05.96 |
| **6** | **rDP** | **02.22** | 6 | CoMMiT-human | 04.50 |
| 7 | CoMMiT-human | 01.36 | 7 | Yang-Server | 04.26 |
| 8 | AIchemy_RNA | 01.03 | 8 | CoMMiT-server | 04.08 |
| 9 | UltraFold | 01.01 | **9** | **rDP** | **04.03** |
| 10 | CoMMiT-server | 00.94 | 10 | UltraFold | 03.63 |
| 11 | Kiharalab | 00.80 | 11 | GWxraylab | 03.54 |
| 12 | Yang | -00.17 | 12 | SoutheRNA | 03.14 |
| 13 | SoutheRNA | -00.37 | 13 | Kiharalab | 03.01 |
| 14 | SHT | -00.45 | 14 | Yang | 02.96 |
| 15 | GWxraylab | -00.66 | 15 | DF_RNA | 02.88 |
| 16 | UltraFold_Server | -00.91 | 16 | LCBio | 02.54 |
| 17 | Coqualia | -01.41 | 17 | Coqualia | 02.35 |
| 18 | GinobiFold | -01.64 | 18 | Rookie | 02.35 |
| 19 | Rookie | -02.58 | 19 | Manifold | 02.30 |
| 20 | Manifold-E | -03.61 | 20 | SHT | 02.23 |
| 21 | Yang- Multimer | -03.75 | 21 | AIchemy_LIG | 02.19 |
| 22 | Manifold | -03.86 | 21 | AIchemy_LIG3 | 02.19 |
| 23 | LCBio | -04.42 | 21 | AIchemy_LIG2 | 02.19 |
| 24 | BAKER | -04.56 | 24 | Yang- Multimer | 02.17 |
| 25 | DF_RNA | -07.60 | 25 | Venclovas | 02.11 |
| 26 | FoldEver | -12.15 | 26 | GinobiFold | 01.81 |
| 27 | Schug_Lab | -12.49 | 27 | UltraFold_Server | 01.79 |
| 28 | FoldEver-Hybrid | -13.39 | 28 | BAKER | 01.63 |
| 29 | Kiharalab_Server | -13.87 | 29 | Manifold-E | 01.56 |
| 30 | CoDock | -14.93 | 30 | PerezLab_Gators | 01.41 |
| 31 | Graphen_Medical | -15.42 | 31 | CoDock | 01.14 |
| 32 | PerezLab_Gators | -16.16 | 32 | Kiharalab_Server | 01.08 |
| 33 | nucE2E | -16.17 | 33 | WL_team | 00.73 |
| 34 | Venclovas | -17.89 | 34 | Schug_Lab | 00.48 |
| 35 | AIchemy_LIG | -19.81 | 35 | nucE2E | 00.39 |
| 35 | AIchemy_LIG3 | -19.81 | 36 | Manifold-LC-E | 00.00 |
| 35 | AIchemy_LIG2 | -19.81 | 36 | UNRES | 00.00 |
| 38 | WL_team | -21.27 | 36 | Manifold-LC | 00.00 |
| 39 | Manifold-LC | -22.16 | 36 | FoldEver | 00.00 |
| 40 | Manifold-LC-E | -22.19 | 36 | FoldEver-Hybrid | 00.00 |
| 41 | UNRES | -22.68 | 36 | Graphen_Medical | 00.00 |

### Supplementary Figures

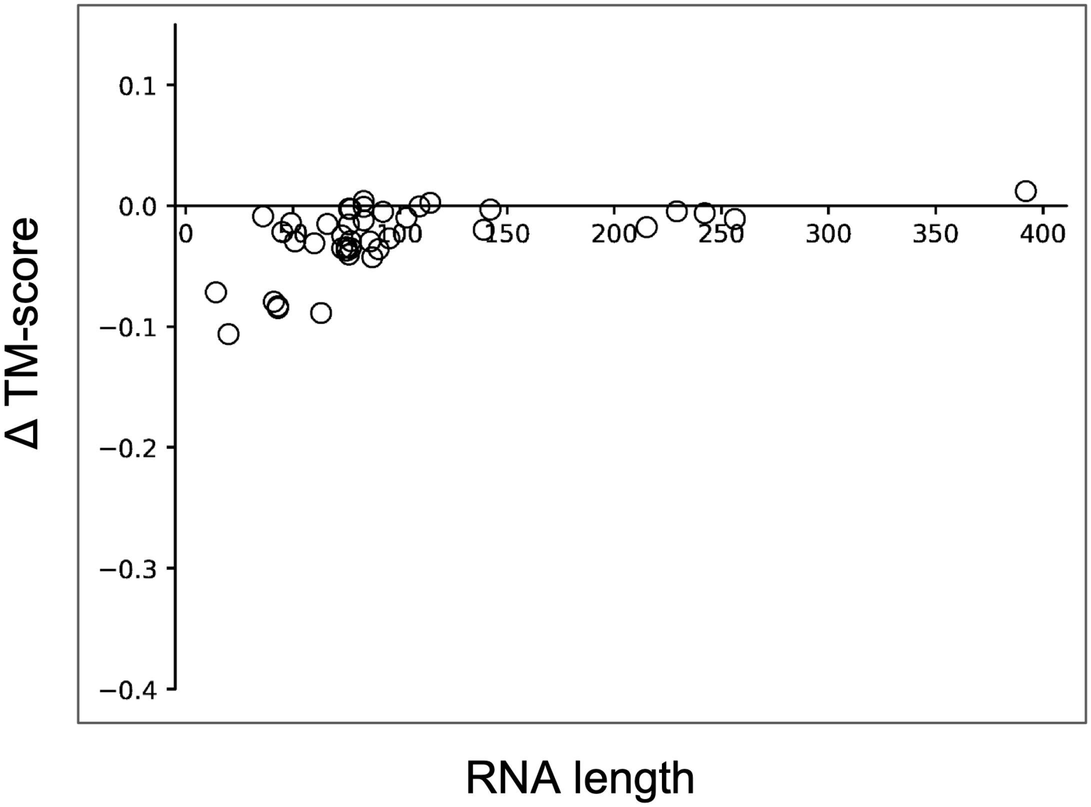


**Figure S1. The difference of TM-score versus the RNA length without geometry potentials, compared to the full pipeline (negative values indicate worse performance for the reduced pipeline).**

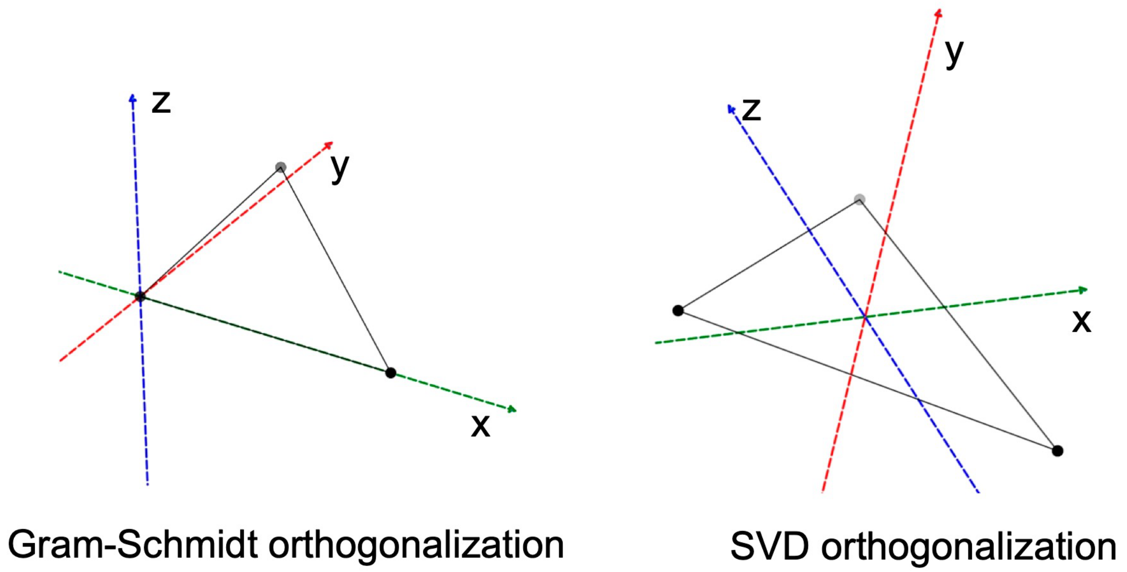


**Figure S2. An illustration of difference between Gram-Schmidt and SVD orthogonalization.**
